# Systematic characterization of short intronic splicing-regulatory elements

**DOI:** 10.1101/2021.05.17.444464

**Authors:** Yuan Gao, Kuan-Ting Lin, Yang Yang, Jialin Bai, Li Wang, Junjie Sun, Lei Sheng, Adrian R. Krainer, Yimin Hua

**Author notes:** To whom correspondence should be addressed at: Jiangsu Key Laboratory for Molecular and Medical Biotechnology, College of Life Sciences, Nanjing Normal University, Nanjing 210023, China, or Cold Spring Harbor Laboratory, PO Box 100, Cold Spring Harbor, New York 11724, USA.

## Abstract

Intronic splicing enhancers and silencers (ISEs and ISSs) are two groups of splicing-regulatory elements (SREs) that play critical roles in determining splice-site selection, particularly for alternatively spliced introns or exons. SREs are often short motifs; their mutation or dysregulation of their cognate proteins frequently causes aberrant splicing and results in disease. To date, however, knowledge about SRE sequences and how they regulate splicing remains limited. Here, using an *SMN2* minigene, we generated a complete pentamer-sequence library that comprises all possible combinations of 5 nucleotides in intron 7, at a fixed site downstream of the 5′ splice site. We systematically analyzed the effects of all 1023 mutant pentamers on exon 7 splicing, in comparison to the wild-type minigene, in HEK293 cells. Our data show that the majority of pentamers significantly affect exon 7 splicing: 584 of them are stimulatory and 230 are inhibitory. To identify actual SREs, we utilized a motif set enrichment analysis (MSEA), from which we identified groups of stimulatory and inhibitory SRE motifs. We experimentally validated several strong SREs in *SMN1*/*2* and *MAPT* minigene settings. Our results provide a valuable resource for understanding how short RNA sequences regulate splicing. Many novel SREs can be explored further to elucidate their mechanism of action.

## INTRODUCTION

Pre-mRNA splicing is an essential step for expression of most eukaryotic genes, during which introns are removed and exons joined to generate a mature mRNA. Exons and introns are either constitutively or alternatively spliced. Alternative selection of 5′ or 3′ splice sites, i.e., alternative splicing, to generate two or more mRNA isoforms is a common phenomenon for pre-mRNAs transcribed from human genes. Natural alternative splicing not only contributes to expansion of transcript diversity, but also serves as a posttranscriptional mechanism to regulate gene function, often in a cell-type- or developmental-stage-specific manner (1). The splicing pattern of a gene in a specific cell setting generally reflects an intricate interplay among multiple cis-acting elements and trans-acting factors, and is influenced by other cellular pathways, such as transcription through chromatin, and signaling pathways.

The strength of four core splicing signals—the 5′ splice site, the 3′ splice site, the poly-pyrimidine tract, and the branch point site—which comprise the first layer of the “splicing code”, is the major determinant of intron and exon definition. However, the prevalence of pseudo-exons with core signals that also match the consensus elements indicates that additional cis-acting splicing signals are involved in distinguishing true exons from pseudo-exons (2,3). These auxiliary signals, termed splicing-regulatory elements (SREs) represent another essential aspect of the splicing code. Based on their role and position, SREs are classified as exonic and intronic splicing enhancers (ESEs and ISEs) or silencers (ESSs and ISSs). Cis-acting SREs are typically located in the vicinity of the splice sites to exert their effects; however, distal elements located >500 nt away from exons, i.e., deep intronic elements, may still affect splice-site selection (4).

SREs can be RNA secondary structures that inhibit splicing by concealing key splicing sequences in double-stranded regions, or in some cases promote splicing by bringing the 5′ and 3′ splice sites in close proximity, or distal enhancer sequences close to their regulated exons (4,5). In most described cases, SREs are single-stranded RNA sequence motifs that are specifically bound by their cognate RNA-binding proteins (RBPs), which function as either splicing activators or repressors. The differences in the splicing pattern of a pre-mRNA in different cell types or developmental stages are mostly due to expression, localization, or phosphorylation alterations of various regulatory splicing factors, which constitute another layer of the regulatory splicing code.

It has long been established that purine-rich and AC-rich ESEs are bound by serine/arginine-rich (SR) proteins (6,7), which facilitate the recognition of adjacent splice sites by the basal splicing machinery. Such activities of SR proteins are often antagonized by hnRNP proteins that recognize splicing silencer sequences (8,9). Recently, an increasing number of RBPs have been found to be involved in splicing regulation. Humans have an estimated 1542 RBPs, or ~7.5% of all protein-coding genes; among them, 692 are mRNA-binding proteins (10). RBPs possess at least one RNA-binding domain, such as an RNA-recognition motif (RRM), a K-homology (KH) domain, an arginine/glycine-rich domain, a DEAD-box motif, or a zinc-finger domain (10,11), and they typically bind to sequence motifs of 3-7 nt (12,13). One characteristic feature of many RBPs is the degeneracy of the sequence motifs they recognize. For example, hnRNP A1, a strong splicing repressor, binds tightly to the consensus pentamer motif UAGGG (13,14). However, the core sequence sufficient for recognition and binding is the dinucleotide AG, with improved affinity for sequences containing UAG or its weaker version, CAG (15). The degenerate nature of RBP-binding motifs makes it challenging to identify authentic binding sites for certain RBPs.

For a known RBP that regulates splicing, in vitro SELEX (systematic evolution of ligands by exponential enrichment) (16) can be employed in conjunction with a splicing assay to identify its functional consensus motif; with this method, functional sequence motifs specific for a subset of SR proteins were uncovered (17). Another frequently used method is CLIP (crosslinking and immunoprecipitation), in which a transcriptome-wide analysis is performed to detect direct protein-RNA interactions in cells; this powerful method has defined a cohort of SRE-RBP pairs that regulate numerous alternative splicing events essential for maintaining cellular homeostasis and function (12,18). With advances in next-generation sequencing technology, transcriptome-wide RNA sequencing is now preferentially used to detect splicing changes and associated SREs regulated by an RBP, whose expression is manipulated via knockdown or overexpression; this approach requires appropriate bioinformatics tools to quantitate isoform reads associated with splicing events (19). On the other hand, to search novel SREs—regardless of the RBPs that recognize them—several strategies have been employed, including conventional deletion and mutational analysis, as well as computational prediction methods, such as RESCUE-ESE (6) and mCross via analysis of published CLIP data (20). Several groups have built minigene reporter systems to screen random-sequence libraries inserted into an alternatively spliced exon or a flanking intron, generating a large amount of information about sequence motifs that influence splice-site selection (21–27). However, despite all the efforts undertaken during the past three decades, our knowledge about SREs—an essential part of the splicing code—remains incomplete.

To better understand short SREs, we generated and analyzed a complete library of pentameric sequences, and directly compared all 1024 sequences inserted at an intronic region in a minigene system. Based on a prior systematic analysis of 207 RBPs from different species, including 85 human proteins, with a method called “RNAcompete”, we know that a large number of RBPs recognize 5-nt or shorter motifs (13). Therefore, pentameric or shorter motifs represent a major fraction of SREs, and they are also at the practical limit of the kind of exhaustive one-by-one analysis we employed here. We chose the spinal muscular atrophy (SMA)-associated *SMN2* as the model gene, because alternative splicing of its exon 7 has been extensively characterized. We chose the mutation region at positions 11 to 15 (CAGCA) in intron 7, a region that is critical for modulation of exon 7 splicing (15,28). The percentage of *SMN2* exon 7 inclusion in multiple cell lines, including HEK293 cells is about 40%, which is ideal to observe changes in exon 7 inclusion in either direction. We analyzed the splicing pattern of each of the 1023 mutants in HEK293 cells, compared to the wild-type (WT) *SMN2* minigene. Our data provide a thorough picture of how different pentameric intronic sequences can regulate an alternative splicing event.

## MATERIALS AND METHODS

### Plasmids

*SMN1*/*2* minigene constructs were pCI-SMN1 and pCI-SMN2 (15). These two minigenes comprise the 111-nt exon 6, a 200-nt shortened intron 6, the 54-nt exon 7, the 444-nt intron 7, and the first 75 nt of exon 8, followed by a consensus 5′ splice site. To obtain all 1023 *SMN2* mutants for making the full pentamer library, we first set up 16 sequence groups in which the first two nucleotides of pentamers were preset (AA, AC, AG, AT, CA, CC, CG, CT, GA, GC, GG, GT, TA, TC, TG and TT) and the remaining three nucleotides were generated by site-directed mutagenesis using a pair of partially-overlapping primers comprising a 3-nt random-sequence pool. 16 pairs of primers were accordingly designed to obtain all mutant plasmids.

A *MAPT* minigene including the 3’-end 256-bp sequence of exon 9, a shortened 388-nt intron 9, the 93-bp exon 10, a shortened 429-nt intron 10, and the 82-nt exon 11 of the *MAPT* gene was constructed in the pCI-neo vector (Promega).

### Cell culture and transfection

HEK293 cells were cultured in Dulbecco’s modified Eagle’s medium (DMEM, Invitrogen) supplemented with 10% (v/v) fetal bovine serum (FBS) and antibiotics (100 U/ml penicillin and 100 μg/ml streptomycin). For splicing analysis, cells were seeded at 1.8×10^5^ per well in 12-well plates; the next day, 1 μg of each minigene plasmid was delivered to cells using branched polyethylenimine reagent (Sigma) in FBS-free DMEM. After incubation at 37 °C for ~6 hours, the transfection solution was removed and replaced with complete medium. Cells were collected for extraction of total RNA 24-36 hrs post-transfection.

### Fluorescence-labeled RT-PCR

Total RNA was isolated from cells with Trizol reagent (Tiangen); 1 μg of each RNA sample was used for first-strand cDNA synthesis with oligo (dT)_18_ and M-MLV reverse transcriptase (Vazyme) in a 20-μl reaction. Transcripts expressed from the *SMN1*/*2* WT and mutant minigenes were amplified semi-quantitatively using 26 PCR cycles with forward primer T7-F2 (5′-TACTTAATACGACTCACTATAGGCTAGCCTCG-3′) and Cy5-labeled reverse primer Ex8-29to52-R (5′-Cy5-TCTGATCGTTTCTTTAGTGGTG TC-3′). Transcripts expressed from the *MAPT* WT and mutant minigenes were amplified using 28-30 PCR cycles with forward primer (5′-CCACTCCCAGTTC AATTACAGC-3′) and reverse primer (5′-Cy5-TAATGAGCCACACTTGGAGGTC-3′). Cy5-labeled PCR products were separated on 6% native polyacrylamide gels, followed by fluorescence imaging with G:Box Chem XL (Syngene), and signals were quantitated by GeneTools software (Syngene). Exon inclusion was expressed as a percentage of the total amount of spliced mRNA.

### Motif Set Enrichment Analysis (MSEA)

For MSEA, a motif set is a group of pentamers with the same sequence feature. Six layers of motif sets were used to describe the sequence features of a pentamer. The first layer uses non-positional motifs, which group pentamers based on 3-mer or 4-mer motifs regardless of the position of the motif (e.g., CCAUG and CAUGC pentamers both belong to the “AUG” motif set). The 2^nd^, 3^rd^, and 4^th^ layers use position-dependent motifs, based on the first (Position 1), the second (Position 2), or the third (Position 3) 3-mers in a given pentamer (e.g., AUGCC and AUGGC pentamers both belong to Position-1 “AUGNN” motif set; N = A, C, G, or U). The 5^th^ layer describes the pentamers based on repeat dimers or single nucleotide repeats (e.g., U:U). It counts how many repeating single or dinucleotides in a given pentamer (e.g., UU is repeated twice in the AUUUU pentamer and is written as UU:UU). The 6^th^ layer is based on the interaction of two nucleotides in the pentamer. For example, we use A-U-- to describe pentamers with an A as the first nucleotide and a U as the third nucleotide. Finally, we used the percent spliced-in (PSI) values of the 1,024 pentamers in two independent replicate experiments to generate the initial data matrix (4×1024) in .gct format. We wrote in-house Perl scripts to generate the motif sets in .gmt format, and generated a ranked list of 1,024 pentamers based on average ΔPSI (mutant – WT) values in .rnk file format. We then used the .gmt and .rnk files for MSEA software (v4.0.3) to compute normalized enrichment score (NES), FDR q-value, and leading edge values (29). Motif sets with FDR q-value < 0.05 and list value < 50% in the leading-edge analysis were considered significant.

### Statistical analysis

Experimental data other than the mutant library data are presented as mean ± standard deviation. Statistical significance was analyzed by Student’s t-test and one-way ANOVA using software SPSS 16.0; a value of *P* < 0.05 was considered statistically significant.

## RESULTS

### Construction of a pentamer library to identify short SREs

To systematically explore pentameric and shorter sequences that are critical for splicing regulation, we took advantage of a previously reported *SMN2* minigene (30) to generate a complete pentamer library by site-directed mutagenesis, and analyzed the effects of all possible mutants on exon 7 splicing. We mutated the 5-nt sequence CAGCA at positions 11-15 in *SMN2* intron 7 into all possible combinations of sequences. We chose this intronic position on the basis of our previous mutational analysis, which showed that suboptimal SREs in this region can noticeably affect exon 7 splicing (15). We obtained a total of 1023 pentamer mutants (**Figure 1A**). We analyzed the splicing patterns of all 1024 WT and mutant minigenes after transient transfection into HEK293 cells, by fluorescence-labeled semi-quantitative RT-PCR. For each experiment, we used the WT *SMN1* and *SMN2* minigenes as controls. We then calculated the difference in the percentage of exon 7 inclusion, i.e., the ΔPSI value, between each mutant and the WT *SMN2* control in two independent experiments.

**Figure 1.**
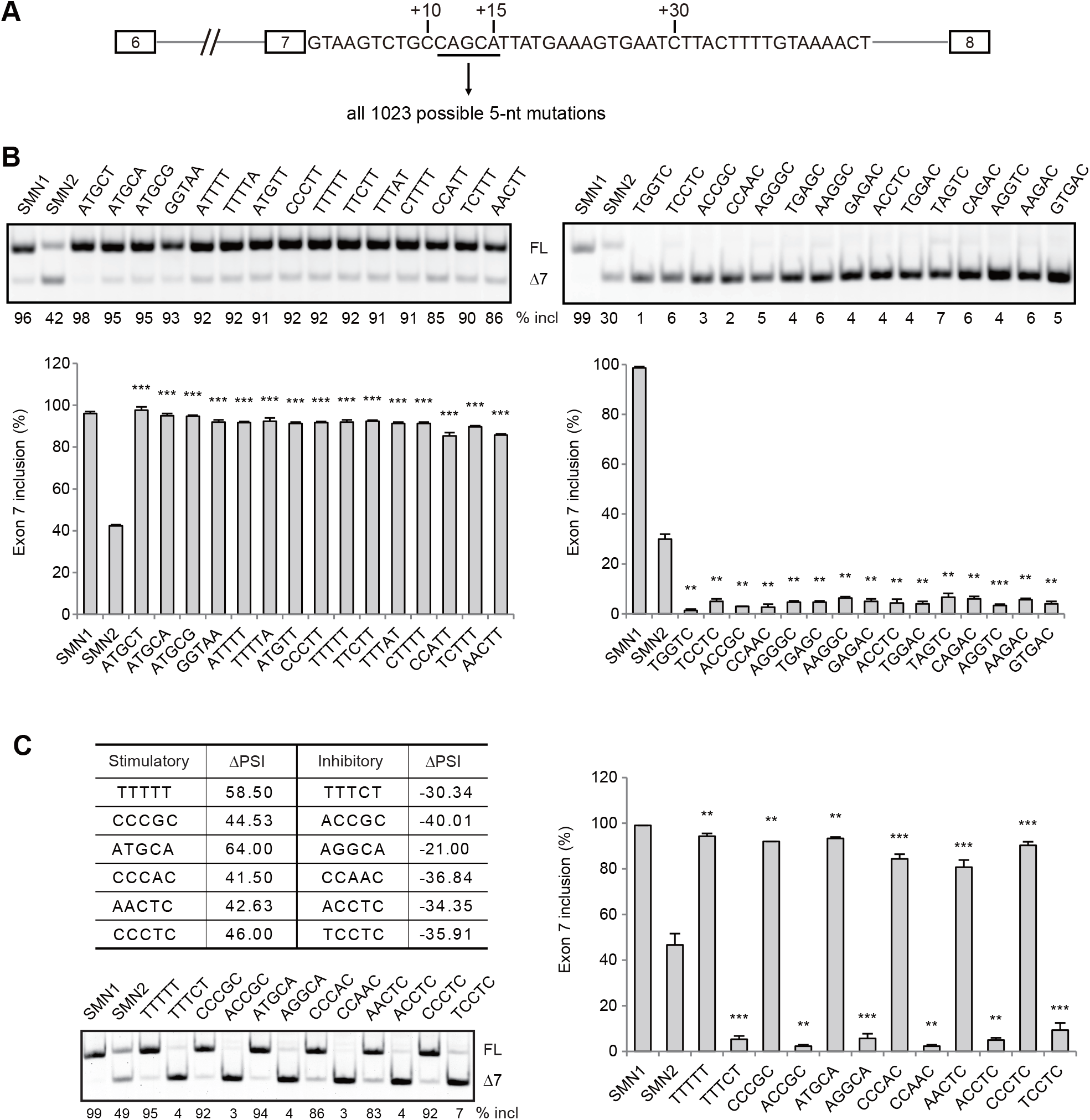
Generation of a complete pentamer-sequence library in an *SMN2* minigene. **(A)** Diagram of the minigene and the insertion site for all the library pentamers. **(B)** Validation of 15 stimulatory pentamers that potently promote exon 7 splicing, and 15 inhibitory pentamers that potently repress exon 7 splicing. **(C)** Large variations in exon 7 splicing are seen for some pentamers with single nucleotide alterations; six pairs were re-evaluated. All WT and mutant minigenes were assayed by transient transfection into HEK293 cells. Quantitative data are shown in histograms (*n* = 3). (**) *P* < 0.01, (***) *P* < 0.001 compared to WT *SMN2*.

### Characterization of the pentamer library with MSEA

PSI values (0~100%) from the pentamer library were highly reproducible in two biological replicates (Pearson’s r = 0.9687, linear regression R^2^ = 0.9310, **Supplementary Figure S1**). The PSI difference between two replicates of WT minigenes ranged from +10% to −21% (−2.52% on average), and the PSI difference between mutant minigenes ranged from +26.48% to −31.84% (+1.23% on average). 686 pentamers had a positive ΔPSI value (up to +64%), and 330 pentamers had a negative ΔPSI value (up to −42%). Seven mutant pentamers had zero ΔPSI (e.g., CUACC). The range of ΔPSI values reflects the fact that the *SMN2* minigene has a base average PSI value of ~41%. 584 pentamers induced >10% exon-inclusion increases, whereas 230 pentamers induced >10% exon-skipping increases (**Supplementary Table S1**). In other words, there were more stimulatory pentamers in the complete pentamer library; one reason may be that the WT pentamer CAGCA is a weak hnRNP A1-binding sequence (15).

The top 50 most stimulatory and inhibitory pentamers, along with 3-nt of flanking sequence on each side, are listed in **Figure 2**. We randomly selected 15 of the top 50 from each category, and re-evaluated their splicing patterns in HEK293 cells, and the data from three independent experiments were consistent with the initial screening (**Figure 1B**). Although the effects of many stimulatory or inhibitory mutants can be attributed to known mechanisms, many of the mutants with strong splicing-regulatory effects represent cis-elements that have not been previously reported. In addition, we observed in many cases that a single nucleotide difference between two mutants led to robust changes in exon 7 inclusion, from being one of the most stimulatory to one of most inhibitory, or vice versa (**Figure 1C**). For example, CCCGC induces 44.53% more exon 7 inclusion, whereas ACCGC induced 40.01% more exon 7 skipping.

**Figure 2.**
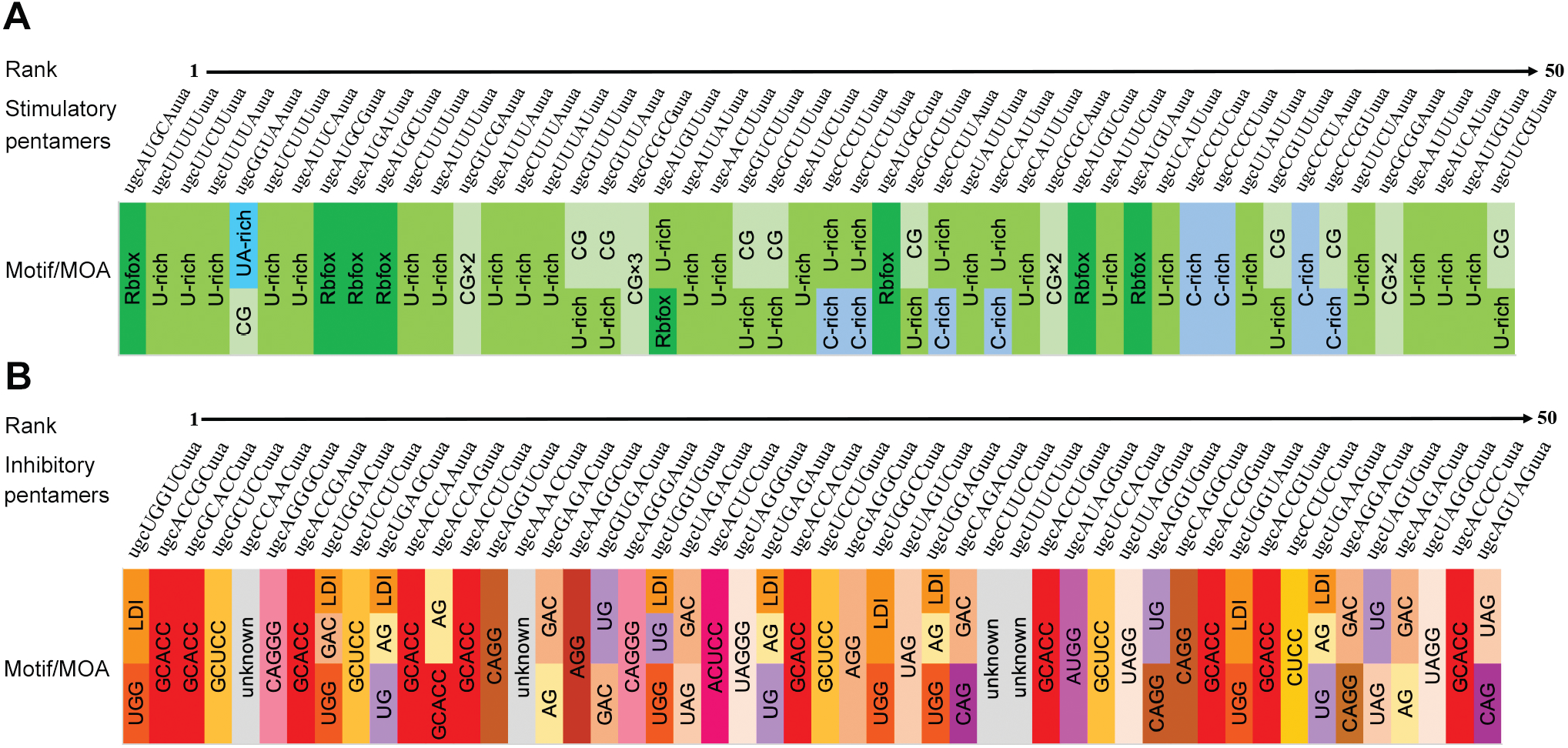
Top 50 most stimulatory and inhibitory pentamers (in uppercase) with flanking sequences (in lowercase), and critical motif(s) or putative mechanism of action (MOA) for each pentamer indicated with a distinct color shade. LDI: enhanced long-distance interaction leading to stabilization of an RNA structure.

Identification of short SREs within or overlapping the stimulatory and inhibitory pentamers should help understand their mechanism of action. To this end, we employed a MSEA approach (see Methods). Initially, 1,216 motif sets were generated based on the six layers of sequence features: (1) Non-positional, (2) Position-1, (3) Position-2, (4) Position-3, (5) Repeat Dimers, and (6) Nucleotide Interaction (**Figure 3A**). Each motif set comprises four to 232 pentamers. To remove extreme cases, we excluded motif sets with fewer than seven pentamers or greater than 200 pentamers. Thus, we used 692 motif sets in MSEA to obtain a normalized enrichment score (NES) based on the ΔPSI ranking of the corresponding pentamers among the 1,024 pentamers in the library (**Figure 3A**). We used the NES score to describe the stimulatory or inhibitory tendency for a given pentamer. A positive NES score means a stimulatory tendency, whereas a negative NES score means an inhibitory tendency. For example, the AUUUU pentamer has all positive NES scores in its six layers of sequence features (**Figure 3A**). This implies a strong stimulatory tendency of the pentamer.

**Figure 3.**
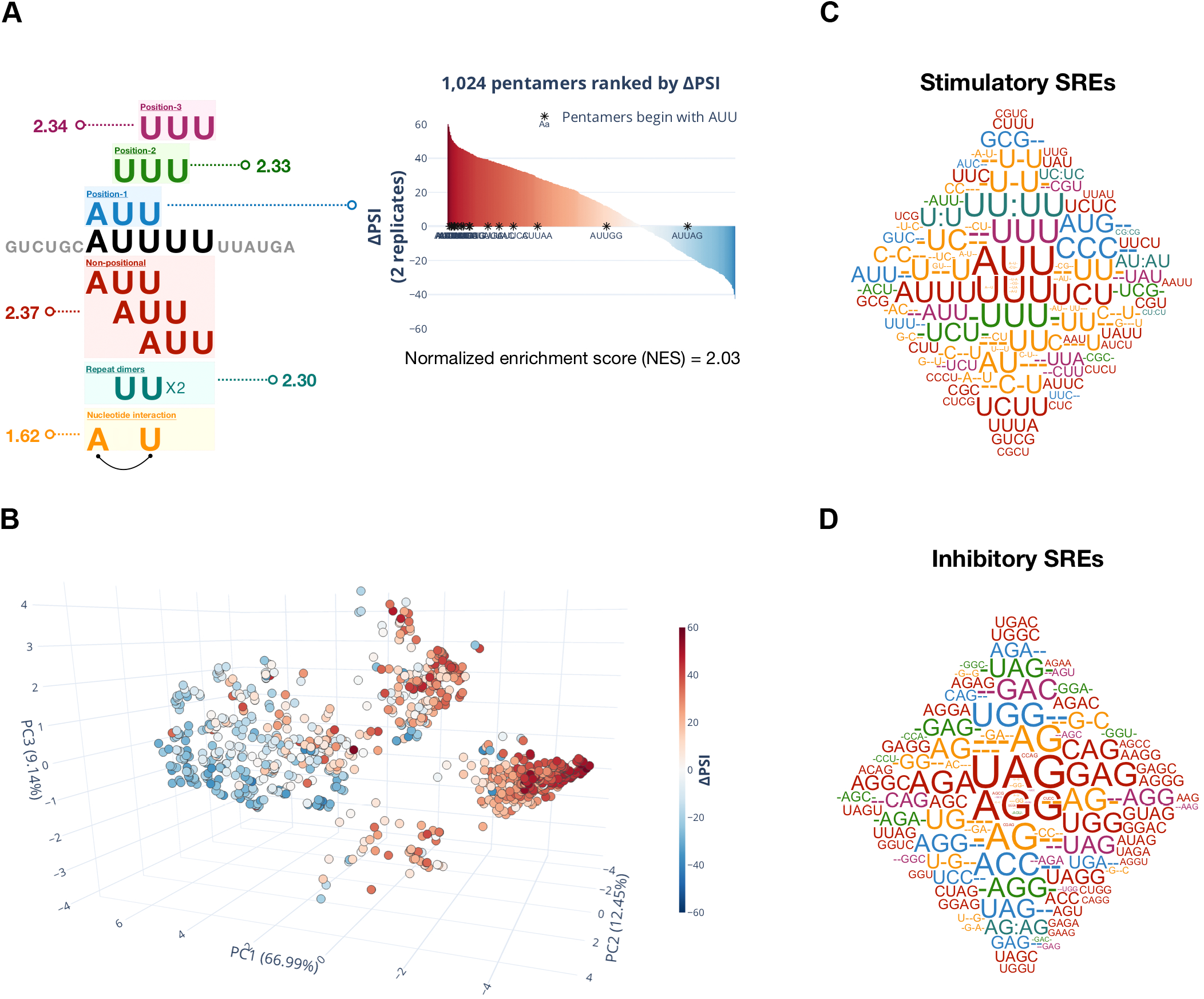
Characterization of the 1,024-pentamer library with MSEA. **(A)** An example pentamer, AUUUU, is shown here to illustrate the six layers of sequence features in MSEA. The non-positional feature can be a 3-mer in any position of the AUUUU pentamer. Position-dependent features (Position 1, Position 2, and Position 3) are based on 3-mers located at a specific position of the pentamer. Repeat-dimer feature is based on a repeating nucleotide(s) in the pentamer. Nucleotide-interaction feature considers the co-existence of any two nucleotides in the pentamer. Each sequence feature has a normalized enrichment score (NES) based on the ΔPSI ranking of pentamers with the same feature in the 1,024-pentamer library. **(B)** A three-dimension PCA plot with 1,024 pentamer balls is shown. Percent variance explained is labeled next to the principal components (PC1, PC2, and PC3). The pentamer balls are colored based on their ΔPSI values (red = exon inclusion; blue = exon skipping). **(C, D)** Stimulatory and inhibitory motifs are shown in C and D, respectively. The sizes of SREs are scaled based on NES scores. SRE motifs are colored based on the layer of sequence feature: Non-positional (red), Position-1 (blue), Position-2 (green), Position-3 (purple), Repeat-dimer (cyan), and Nucleotide-interaction (yellow).

To project the distance between pentamers in the 6-layer space, we performed Principal Component Analysis (PCA). We used the first three principal components to generate a 3-dimensional PCA plot (**Figure 3B**). The first two principal components (PC1 and PC2) cover 79.44% of the variance. With the additional third principal component (PC3), the PCA plot covers 88.58% of the variance. We observed two major clusters in the PCA plot: (1) a stimulatory cluster (red); and (2) an inhibitory cluster (blue) (**Figure 3B**). This shows that the 6-layer sequence features can nicely separate the stimulatory from the inhibitory pentamers.

Among the non-positional motif sets, 30 stimulatory and 46 inhibitory 3- and 4- mers were significant. Eight stimulatory and nine inhibitory Position-1 3-mers, six stimulatory and 12 inhibitory Position-2 3-mers, as well as seven stimulatory and 14 inhibitory Position-3 3-mers were all significant. Repeat dimers had six stimulatory (including single nucleotide U:U repeats) and one inhibitory (AG:AG) SRE motifs. Nucleotide-interaction motif sets had 38 stimulatory and 23 inhibitory SRE motifs. Overall, stimulatory SRE motifs are more diverse (e.g., UUU, AUG, and CCC), whereas inhibitory SRE motifs have strong preferences for A and G nucleotides. Also, inhibitory SRE motifs in general have higher NES scores (**Figure 3C and D, Supplementary Table S2**).

### Classifications of motifs enriched by non-positional MSEA

Among 76 3- and 4-mers enriched by non-positional MSEA, the two most inhibitory motifs are “UAG” and “AGG”, with normalized enrichment scores (NES) of −4.7678 and −4.4144, respectively (**Figure 4A and B, Supplementary Table S2**). These correspond to the binding motifs of hnRNP A1 (14), whereas the two most stimulatory motifs are “UUU” (NES = 2.4378) and AUU (NES = 2.3903) (**Figure 4C and D, Supplementary Table S2**), which are likely recognized by TIA1 (13,19,31).

**Figure 4.**
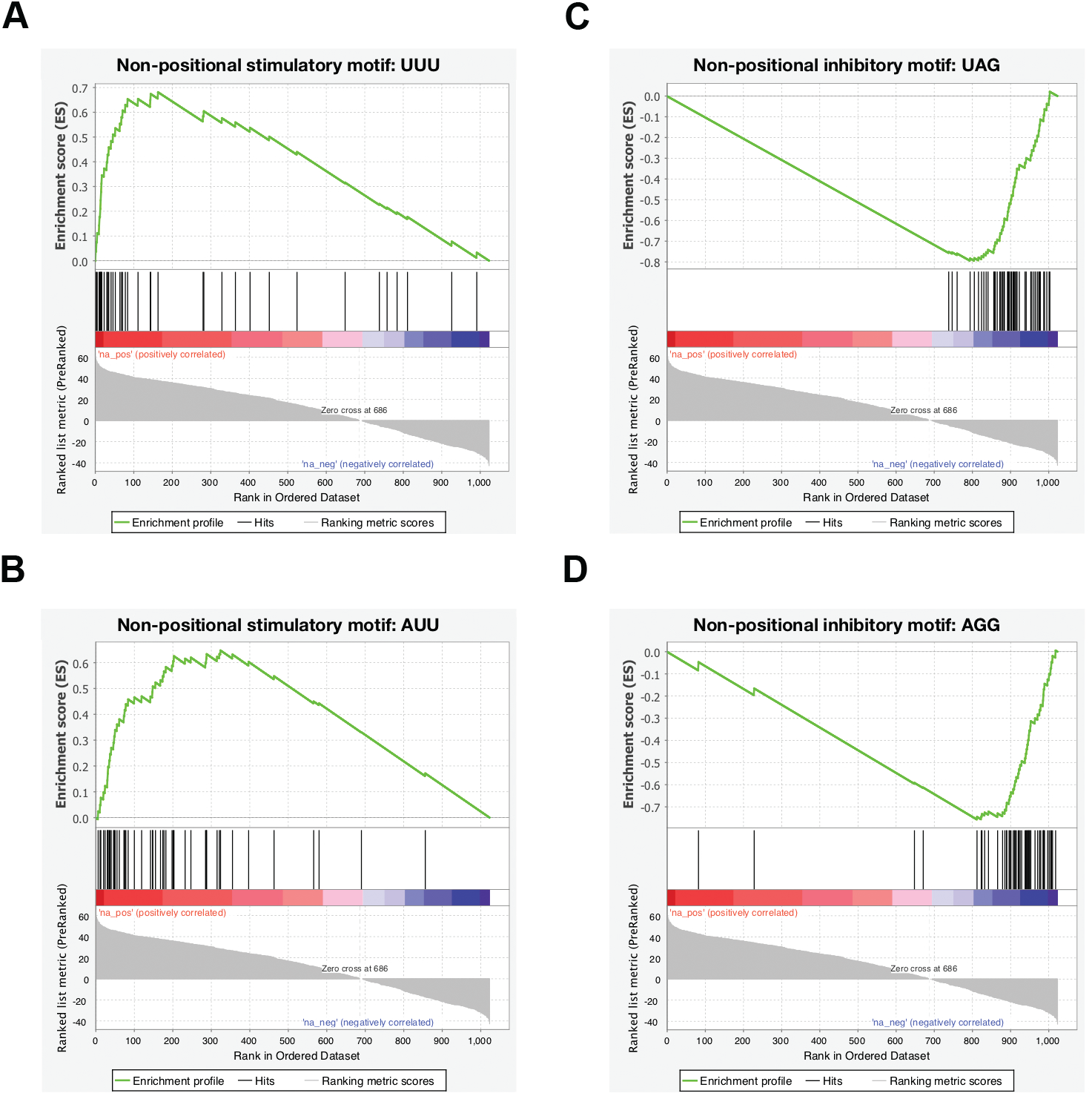
Enrichment plots of top non-positional motifs. **(A and B)** The top two stimulatory motifs, UUU and AUU, are shown to illustrate the ΔPSI rankings of their corresponding pentamers in the 1,024-pentamer library. Red and blue colors in the spectrum in the middle represent pentamers with positive and negative ΔPSI value, respectively. **(C and D)** The top two inhibitory motifs, UAG and AGG, are shown here to illustrate the ΔPSI rankings of their corresponding pentamers in the 1,024-pentamer library. The spectrum in the middle is as in **(A and B)**.

We looked at the sequence features of the 30 enriched stimulatory SRE motifs, and found that they can be separated into two major classes, which we termed U-rich and CG-containing (CG-core), plus one minor class that is C-rich (**Figure 5A**). 19 motifs belong to the U-rich class, with each motif containing at least two Us. Interestingly, only one of the U-rich motifs has a G (UUG). As the immediate tetranucleotide downstream of the library site is UUAU, this suggests that poly-U runs or U-rich sequences (> 4 Us) mixed with scattered A or C tend to be strong ISEs. We interrogated all 32 pentamer mutations that are comprised of solely A and/or U, and found 20 with ΔPSI > 30 and all with a positive ΔPSI, reflecting that UA-rich sequences are generally ISEs. However, pentamers with 3-5 Us have much higher ΔPSI values than those with 3-5 As, pointing to the Us as the key component of these motifs (**Supplementary Table S3**); this explains why AAU is the only enriched motif with more As than Us. On the other hand, U-rich motifs mixed with scattered C, though mostly stimulatory, can be very inhibitory in some cases (**Figure 2, Supplementary Table S4**). It is reasonable to assume that most U-rich motifs function by binding to TIA-1 and TIAL; however, we cannot rule out other mechanisms that may mediate the effects of particular motifs, considering the existence of multiple RNA-binding proteins with high affinity for U-rich sequences (13).

**Figure 5.**
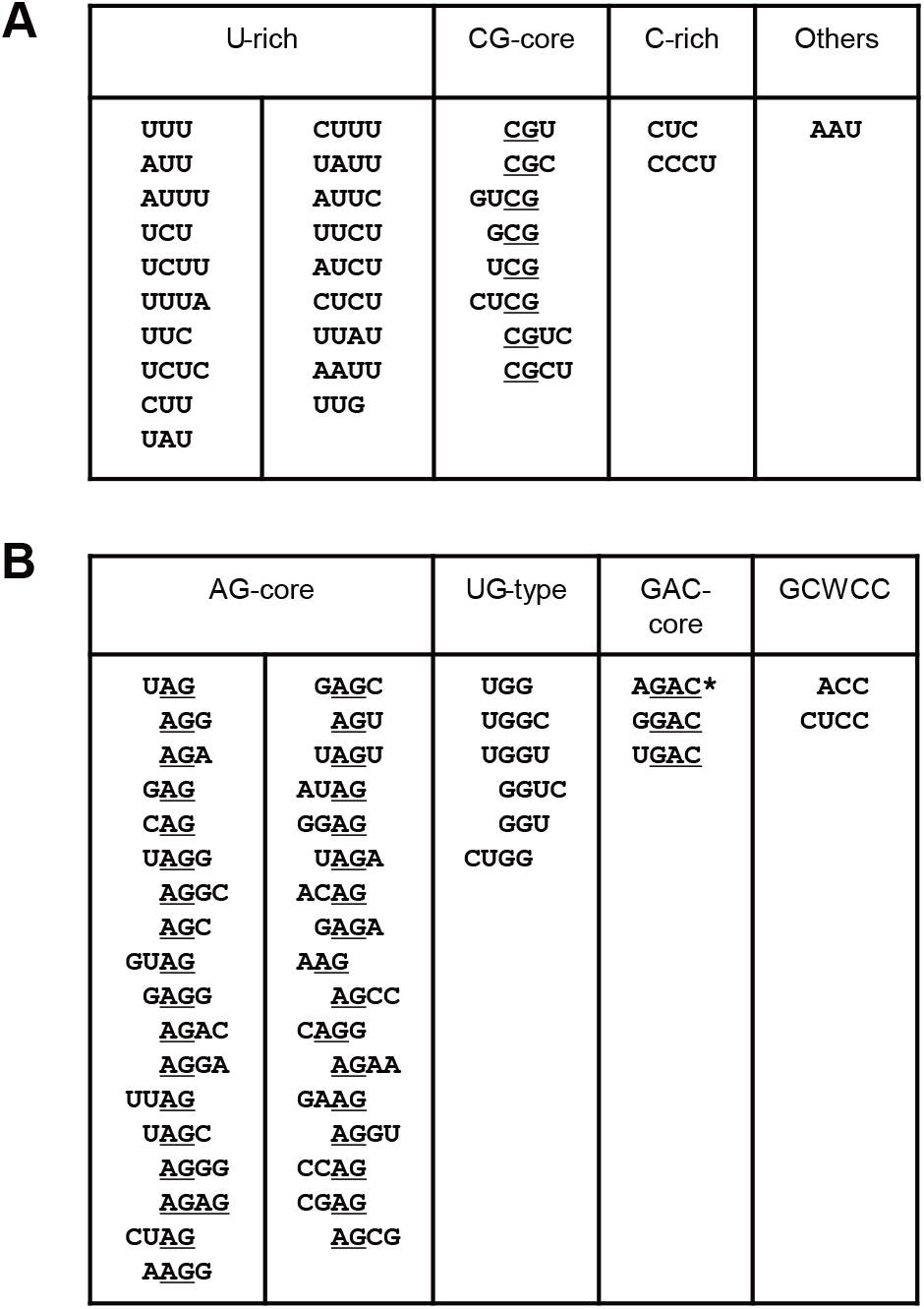
Classifications of 3-4-mer motifs enriched by non-positional MSEA. **(A)** We separated 30 enriched stimulatory motifs into three main groups. U-rich motifs have at least two Us. CG-core motifs have one copy of CG, and C-rich motifs have at least two Cs. AAU supposedly forms a UA-rich splicing enhancer with the 4-nt sequence UUAU immediately downstream of the pentamers. **(B)** We separated 46 enriched inhibitory motifs into four groups: AG-core motifs have at least one copy of AG; UG-type motifs have either UGG or GGU; and GAC-core motifs have the trinucleotide GAC but preclude C as the nucleotide immediately preceding it. The enrichment of ACC and CUCC confirms that the pentamers GCACC and GCUCC (GCWCC) are potent ISSs.

The second major class of stimulatory motifs includes eight motifs (CGU, CGC, GUCG, GCG, UCG, CUCG, CGUC and CGCU), all of which share the dinucleotide CG, highlighting this CG feature as the core of this type of splicing enhancers (**Figure 5A**). Alignment of all the 8 CG-core motifs suggests that UCGY (Y = pyrimidine) is a stronger extended version of the CG dinucleotide.

We designated the remaining two motifs, CUC and CCCU, both with more Cs (at least two Cs) than Us, as C-rich motifs (**Figure 5A**). As the first upstream flanking nucleotide is C, this hints that C-rich sequences, such as three or more consecutive Cs, or mixed with scattered Us, constitute an ISE. We discuss this further below (Position 1-dependent MSEA).

The 46 inhibitory motifs—11 3-mers and 35 4-mers—enriched by non-positional MSEA can be separated into four classes (**Figure 5B**). Among the 11 inhibitory 3-mer motifs, eight contains at least one A and one G, and all display a pattern of NAG or AGN, including all possible combinations, but not BGA (B = C, G, or U) or GAH (H = A, C, or U), confirming that the dinucleotide AG, rather than GA, is the core of purine-rich silencers. Most of the NAG/AGN motifs rank at the top of the 46-motif list (**Supplementary Table S**2). We designated those motifs containing at least one AG as AG-core motifs, which represent the most predominant class of inhibitory motifs and includes 27 4-mers, in addition to the eight 3-mers (**Figure 5B**). Among the 27 4-mer motifs, 16 are NUAG, UAGN, NAGG, and AGGN, consistent with UAGG being the SELEX winner sequence with high affinity for hnRNP A1 (14). The predominant enrichment of AG-containing motifs by non-positional MSEA supports the notion that hnRNP A/B proteins, particularly the abundant hnRNP A1 and hnRNP A2, are among the strongest splicing repressors.

We classified the seven enriched 3- and 4-mers that contain a copy of either UGG or GGU (UGG, UGGC, UGGU, GGUC, GGU, CUGG and UGGA) as UG-type motifs (**Figure 5B**). The silencing activities of these motifs may involve multiple mechanisms. It has been reported that two 8-nt sequences in intron 7, one from position 3 to 10 and the other from 281 to 289, form a double-stranded RNA structure, which impairs the annealing of U1 snRNA to the 5′ splice site of intron 7, contributing to the predominant skipping of *SMN2* exon 7 (32). We found that mutations in either strand that presumably strengthen the long-distance RNA-RNA interaction also repressed exon 7 splicing in the *SMN1* minigene (**Supplementary Figure S2**). UGNNN and UGGNN extend the predicted double-stranded RNA structure by at least two and three base pairs, respectively. Therefore, this may be an important part of the mechanism that makes them strongly inhibitory. On the other hand, UGG and GGU themselves have inhibitory activity, particularly when they are flanked with pyrimidines, as in UGGU, UGGC, CUGG and GGUC, and the consensus motif appears to be UGGY. Indeed, 14/16 NUGGY and UGGYN pentamers in the library strongly inhibited exon 7 splicing compared to the WT *SMN2* minigene (**Supplementary Table S5**); the two exceptions were AUGGU and AUGGC, both of which are part of an RBFOX-binding motif (UGCAUG) (see below). 16 NUGGN pentamer mutants provide an opportunity to analyze the UGG motif itself, with little interference from the RNA secondary structure. When UGG is followed by G or A instead of C or U, its inhibitory activity is markedly reduced or even reversed (**Supplementary Table S6)**. This observation suggests that GGG and GGA are at best weakly inhibitory, whereas GGG may be even slightly stimulatory in some cases, and counteracts UGG. It has been established that intronic G-rich motifs or poly-G runs act as splicing enhancers (33,34). Therefore, we interrogated all 40 pentamer mutants with a G triplet (NNGGG, NGGGN, and GGGNN). Interestingly, except for seven AGG- and three UGG-containing pentamers, all others have a ΔPSI > 0 and 18 have a ΔPSI > 17 (**Supplementary Table S7**). For all pentamers containing four consecutive Gs, 5/7 robustly promoted exon 7 inclusion, the exceptions being AGGGG and UGGGG (**Supplementary Table S7**), confirming that poly-G stretches are indeed ISEs.

We classified three 4-mers (AGAC, GGAC and UGAC) as GAC-core motifs (**Figure 5B**). AGAC also falls into the AG-core class. Interestingly, when the nucleotide preceding GAC is C, its inhibitory activity is neutralized. We looked at all mutants containing CGAC, and found that 18/24 markedly promoted exon 7 splicing, compared to the WT *SMN2* minigene with ΔPSI > 11 (**Supplementary Table S8**). On the other hand, 22/24 mutants that comprise DGAC (D = A, U or G) had a ΔPSI < 0, and 18 markedly inhibited exon 7 splicing (ΔPSI < 12) (**Supplementary Table S9**). These data support DGAC motifs as a class of ISSs.

ACC and CUCC, which rank No. 15 and No. 45, respectively, on the enriched inhibitory motif list, are part of two pentamers, GCACC and GCUCC; both mutants potently inhibited exon 7 splicing, with ΔPSIs of −38.50 and −37.50, respectively (**Figure 2**, **Supplementary Table S1**). Here we named these elements GCWCC-type (W = A or U) (**Figure 5B**).

### Motifs enriched by position-dependent MSEA

Non-positional analysis generally identifies fully functional motifs in the library, but may miss motifs that span one of the two junctions; such motifs require position-dependent analysis to pinpoint. For example, some motifs may include the upstream trinucleotide UGC or the downstream dinucleotide UU, or a portion thereof, to form a fully functional motif.

For Position 1-dependent MSEA, we analyzed all 3-mers at Position 1 (nucleotides 1-3 in each pentamer), which identified 17 3-mers that strongly affected exon 7 splicing, with 8 stimulatory and 9 inhibitory ones (**Figure 6A and B, Supplementary Table S2)**. 4/8 stimulatory motifs (GCG, AUU, UUU and UUC) were already enriched by non-positional MSEA. Three new motifs CCC, AUG, and GUC appear to rely on the upstream flanking nucleotides to form a functional or stronger motif (**Figure 6A**). The immediate upstream nucleotide C and CCC make a longer C-rich motif, and the C and GUC form the CG-core motif CGUC, which is on the enriched non-positional MSEA list. CCC is ranked as the top stimulatory motif on the list, though it failed to show enrichment by non-positional MSEA, indicating that poly-C runs (4 or longer) are one class of strong ISEs.

**Figure 6.**
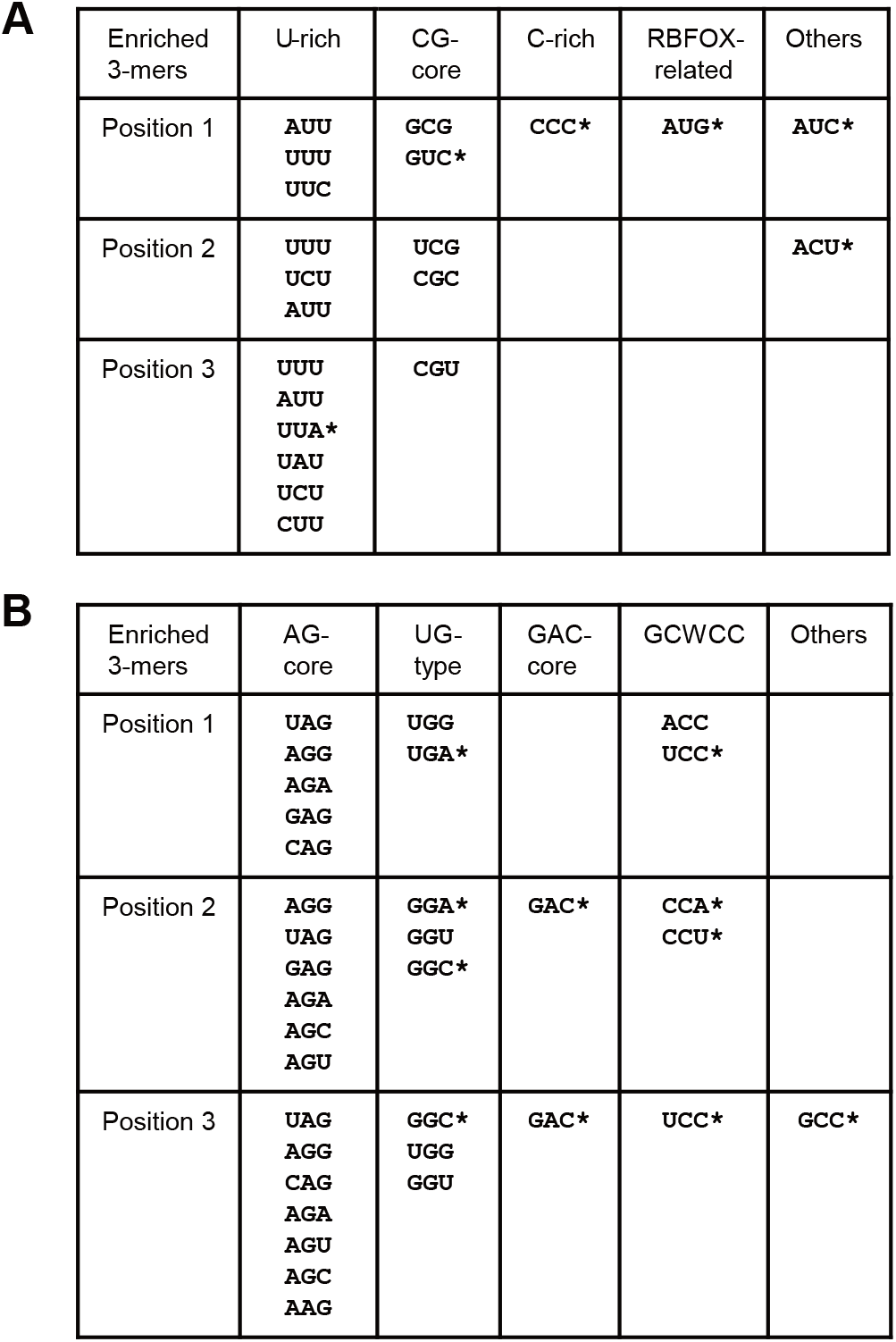
Classification of 3-mer motifs enriched by Position 1-, Position 2-, and Position 3-dependent MSEA. Six additional stimulatory motifs **(A)** and 11 additional inhibitory motifs **(B)** (marked with *) were identified by positional MSEA. AUG, CCC and GUC apparently rely on the upstream UGC or C to form UGCAUG, CCCC or CGUC to form a full or stronger ISE, whereas ACC and UCC rely on the upstream GC to form the potent ISSs GCACC and GCUCC. UGA at Position 1 strengthens an RNA secondary structure (**Supplementary Figure S2**), whereas GAC at Position 2 or Position 3 precludes the formation of CGAC in most cases.

It was not surprising to find AUG among the top two on the list, as it is part of a well-known splicing enhancer, UGCAUG, with the upstream flanking trinucleotide UGC (**Figure 6A**). UGCAUG is the binding motif of the RBFOX family of proteins. It has been well documented that these proteins promote splicing when binding downstream of the 5′ splice site, including *SMN2* intron 7 (35,36). We classified these motifs as RBFOX-related. AUC appears not to rely on upstream sequences to form a stimulatory motif, and its enrichment at Position 1 is likely due to other 3-mers at this position forming inhibitory motifs with the upstream GC (GCWCC) or C (CAGN).

Regarding the 9 inhibitory 3-mers enriched by Position 1-dependent MSEA (**Supplementary Table S2**), UGG and ACC replaced UAG and AGG as the top 2 inhibitory motifs, compared to the non-positional analysis, suggesting that upstream flanking nucleotides contribute to their silencing activities. Two 3-mers (UCC and UGA) are new. UGG and UGA can be explained by the above-mentioned long-distance RNA secondary structure. On the other hand, with the upstream dinucleotide GC, ACC and UCC form GCACC and GCUCC, respectively, which further confirms GCWCC as a class of potent splicing silencers (**Figure 6B**).

For Position 2-dependent MSEA, we analyzed all 3-mers at Position 2 (nucleotides 2-4 in each pentamer), which led to the identification of six stimulatory and 12 inhibitory motifs (**Figure 6A and B**, **Supplementary Table S2**). 5/6 stimulatory motifs, U-rich or CG-core, were enriched by non-positional MSEA, except ACU, which similar to AUC identified by Position 1-dependent MSEA, appears not to rely on flanking sequences to form a functional ISE, but rather avoids the formation of silencers (such as AG, DGAC, UGG and CUCC) when positioned at 2-4. Among the 12 enriched inhibitory motifs, five (GAC, GGC, CCU, CCA and GGA) are new, but they all belong to the above-defined motif classes (**Figure 6B**). Six are AG-core and all were enriched by non-positional MSEA. That GAC is enriched here is most likely because its inhibitory activity is abolished or compromised when placed at Position 1, due to the upstream C. GGC, GGU and GGA apparently rely on the first nucleotide in the pentamers to be A or U, to form AG-core or UG-type motifs, whereas CCU and CCA rely on the first nucleotide to be A or U to make GCWCC silencers.

For Position 3-dependent MSEA, we analyzed all 3-mers at Position 3 (nucleotides 3-5), which identified seven stimulatory and 14 inhibitory 3-mers (**Figure 6A and B**, **Supplementary Table S2**). Except for UUA, all others are on the stimulatory-element list identified by non-positional MSEA. Together with the downstream dinucleotide UU, UUA forms UUAUU, which is among the top 50 stimulatory pentamers in the library (**Figure 2**). Another contributing factor is that it avoids the formation of the strong inhibitory motifs UAG and UAGG at nucleotide positions 3-5. Among the 14 inhibitory 3-mers, only GCC was not enriched above. We examined the 16 NNGCC mutants, but only half were inhibitory, and they include five comprising the AG-core, two comprising UGG motifs, as well as GCGCC, which is likely a weak version of GCWCC (**Supplementary Table S1**). Therefore, GCC is likely a moderate or weak ISS motif; its enrichment is partly because G at position 3 allows the formation of UAG, CAG, CAGG, AGG and UGG. This also explains why GAC replaced UAG as the most inhibitory 3-mer at positions 3-5.

### Dinucleotide and nucleotide interaction analyses

Analysis of 3- and 4-mer motifs enriched by MSEA demonstrates that sequences as short as two nucleotides play critical roles in defining SREs, which prompted us to explore whether any two nucleotides can indeed be enriched with significant splicing-regulatory activity by MSEA. Repeat-dimer analysis identified five dinucleotides (UU, AU, UC, CG and CU) as stimulatory, and only one dinucleotide (AG) as inhibitory; single-nucleotide U repeats also displayed stimulatory activity (**Supplementary Table S2**). We believe that UU, AU, UC, and CU heavily rely on the downstream UUAU to form U-rich ISEs, and thus none is sufficient to be a specific dinucleotide core of SREs. The only stimulatory dinucleotide core is CG, whereas the only inhibitory dinucleotide core is AG. As CG and AG constitute opposite splicing signals, a single nucleotide C>A mutation in CG-core motifs is expected to result in severe splicing defects. One such case is associated with breast cancer, in which the deleterious missense mutation c.5242C>A in *BRCA1* causes exon 18 skipping, resulting in the loss of 26 aa that are essential for the protein’s function (37).

We next performed nucleotide-interaction analysis (see MSEA in Methods) and identified 39 stimulatory nucleotide interactions, including eight U/U, 11 U/C, 11 U/A, two C/C, three G/U, one G/C, two CG, and one AC (**Supplementary Table S2**). 30/39 are U/U, U/C and U/A interactions, confirming that poly-U or U-rich sequences with scattered C or A are generally strong ISEs. All four G/U and G/C interactions have the G at nucleotide position 1, and thus the G forms a CG dinucleotide with the upstream C, consistent with CG being the core of a class of ISEs. Indeed, the CG dinucleotide itself was enriched twice on the list of stimulatory nucleotide interactions. We also identified 23 inhibitory nucleotide interactions, which include nine A/G (among them four AG dinucleotides and two GA dinucleotides), four U/G (all Us being at position 1), four G/G, three C/C, two G/C, and one AC at positions 1-2 (**Supplementary Table S2**). The nucleotide-interaction analysis is consistent overall with our classification of stimulatory and inhibitory motifs obtained by positional and non-positional MSEA.

### Poly-C and poly-U tracts are comparable in promoting *SMN2* exon 7 splicing

We next validated several interesting findings from the pentamer-library analysis. Multiple U-rich pentamers, such as UUUUU, UUCUU and UUUUA rank among the strongest stimulatory mutants (**Figure 2**), and form longer U-rich stretches with the downstream nucleotides UUAU. On the other hand, based on the Position 1-dependent MSEA, sequences with four or more consecutive Cs also strongly promote exon 7 inclusion. To gain further insight into splicing regulation by poly-U and poly-C sequences and compare their effects, we inserted different lengths of Us or Cs at two positions: one after position 10 (P10) and one after position 15 (P15) of intron 7 in the *SMN2* minigene. These mutants were analyzed in HEK293 cells. As shown in **Figure 7**, four or more Us inserted at P10 robustly promoted exon 7 splicing, with the percentage of exon 7 inclusion increasing from 48% (WT *SMN2*) to 76% (4 Us) or higher (> 4Us). Insertion of three-Us at P15, which forms a 5-U tract with the 2Us downstream, increased the percentage to 54%. At both positions, the longer the inserted poly-U tract, the greater the exon 7 inclusion. Similarly, when we inserted three Cs at P10, which form a 5-C tract with the two flanking Cs, the percentage of exon 7 inclusion increased from 50% in *SMN2* to 67%, and there was a strong correlation between the number of Cs and exon 7 inclusion (**Figure 7B**). However, when we placed poly Cs at P15, strong suppression of exon 7 splicing ensued, which, we believe, is due to the immediate upstream trinucleotide GCA forming a strong GCACC ISS with the poly-C sequences. Indeed, disruption of the GCACC motif by insertion of an extra A or U before the poly-C stretches restored the stimulatory activity of the poly-C tracts (**Figure 7C**). These data confirm that poly-C sequences have similar effects as poly-U in promoting exon 7 inclusion.

**Figure 7.**
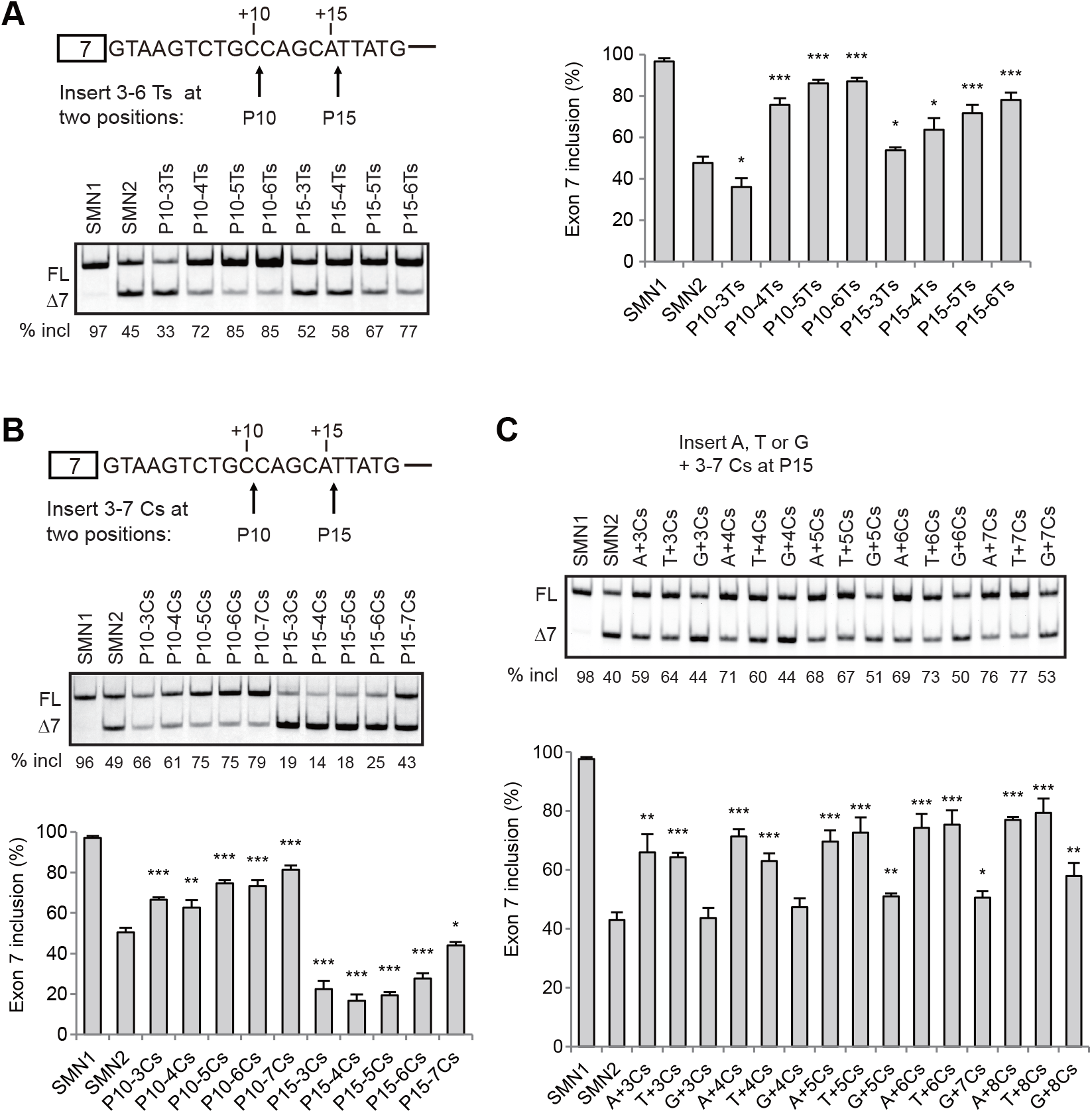
Comparison of the effects of poly-C and poly-U stretches on *SMN2* exon 7 splicing in HEK293 cells. **(A-B)** Insertion of poly-U or poly-C runs at two sites: one after position 10 (P10) and one after position 15 (P15) in *SMN2* intron 7. We named the mutants based on the insertion site and the number of Us or Cs. **(C)** We inserted a single A, U, or G at P15 preceding the poly-C runs in the P15 poly-C mutants, to abolish the formation of GCACC, a novel strong ISS. We named the mutants with the inserted extra nucleotide and the number of Cs; for example, “A+3Cs” means insertion of ACCC. Quantitative data are shown in histograms (*n* = 3). (*) *P* < 0.05, (**) *P* < 0.01, (***) *P* < 0.001 compared to WT *SMN2*.

### GCACC and GCUCC are strong ISSs

Among the top 13 inhibitory pentamers, eight are attributable to GCACC or GCUCC (**Figure 2**), whose silencer activities we confirmed by Position 1-dependent MSEA (**Figure 5B**) and the above poly-C insertion study (**Figure 7**). To further test GCACC as a novel strong ISS motif, we placed it in different sequence settings by inserting it at four positions: P15, P24, P33, and P43 in intron 7 of both the WT *SMN2* and *SMN1* minigenes; insertions at these sites cause no disruptions of known SREs. To avoid the formation of AG and AGG at P24 and P33, we also inserted UGCACC at these two sites. We transfected all insertion mutants into HEK293 cells for splicing analysis. As shown in **Figure 8A**, insertions of GCACC or UGCACC at P15 and P24 robustly repressed exon 7 splicing in *SMN2*, and the effect was less but still pronounced in *SMN1*. The effect of GCACC inserted at P33 and P42 was drastically compromised or abolished, suggesting that close proximity to the 5′ splice site is crucial for this element to exert its silencing activity.

**Figure 8.**
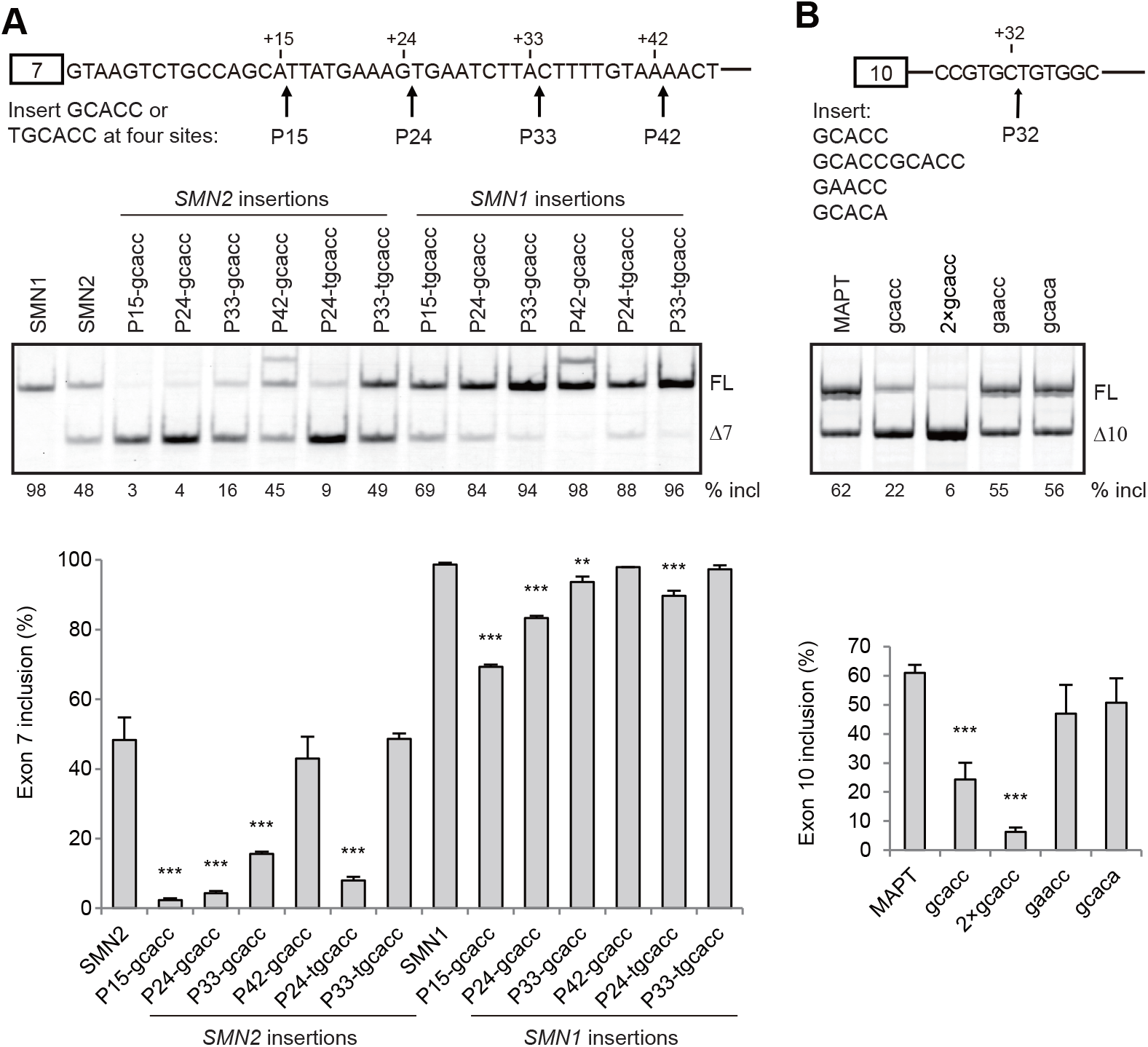
The effects of GCACC on *SMN1/2* exon 7 and *MAPT* exon 10 splicing assayed in HEK293 cells. **(A)** We inserted GCACC at four sites: after positions 15 (P15), 24 (P24), 33 (P33) and 42 (P42), respectively, in both the *SMN1* and *SMN2* minigenes. Insertion of GCACC at P24 and P33 creates an AGG and AG, respectively. Therefore, we also tested insertion of UGCACC at P24 and P33 to evaluate the effects of the newly formed AGG and AG. The data show that the closer the GCACC motif is to the 5′ splice site of intron 7, the stronger its inhibitory effect is on exon 7 splicing. **(B)** We tested one or two copies of GCACC in a *MAPT* minigene by insertion after position 32 (P32) in intron 10. GAACC and GCACA served as controls. Robust inhibition of exon 10 inclusion is seen by GCACC, but not the two controls. The effect of two copies was even stronger. Quantitative data are shown in histograms (*n* = 3). (**) *P* < 0.01, (***) *P* < 0.001 compared to WT *SMN2, SMN1 or MAPT*.

To explore whether GCACC is a general SRE that regulates other alternative splicing events, we tested it in a *MAPT* minigene, which comprises part of exon 9, a truncated intron 9, exon 10, intron 10 and exon 11. Resembling the endogenous gene, exon 10 of the minigene is alternatively spliced (**Figure 8B**). An RNA secondary structure is present at the 5′ splice site of *MAPT* intron 10 and regulates exon 10 splicing (38,39). To avoid disruption of the RNA structure and other natural SREs, we inserted one or two copies of GCACC, following position 32 (P32) of intron 10, with two neutral pentamers (GAACC and GCACA) as controls (**Supplementary Table S1**). We analyzed the WT minigene and the insertion mutants after transfection into HEK293 cells. As predicted, insertion of one copy of GCACC, but not GAACC or GCACA, at P32 markedly reduced exon 10 inclusion from 62% (WT) to 22%; and two copies of GCACC reduced exon 10 inclusion further, to 6% (**Figure 8B**), confirming GCACC as a general and potent ISS motif.

GCACC and GCUCC differ by one nucleotide. Whether their mechanisms of action are similar is unknown. We note that the first G is essential for the strong inhibitory activity of GCACC. In contrast, the first G appears not to be strictly required in GCUCC, as both mutants ACUCC and CCUCC in the pentamer library strongly inhibited exon 7 splicing, though with slightly weaker effects than GCUCC. The exception is mutant UCUCC, which had weak inhibitory activity, with a ΔPSI of −3.50.

### CG repeats potently promote *SMN2* exon 7 splicing

We showed above that CG-containing motifs are a new class of ISEs, of which UCGY is a particularly strong version. Intriguingly, pentamers with two copies of CG tend to be more stimulatory than those with only one copy of CG (**Supplementary Tables S1 and S2**). Indeed, GCGCG, the only mutant that has three CG repeats, counting the upstream C, is one of the top stimulatory pentamers in the library (**Figure 2),** suggesting that the stimulatory activity of each CG copy is additive.

To verify that CG repeats are indeed potent ISE motifs, we inserted CGCGCG at P15, P24, P33 and P42, respectively, in intron 7 of the *SMN2* minigene, and examined exon 7 splicing of these mutants in HEK293 cells. Insertions of CGCGCG at the three proximal sites, but not at P42, robustly promoted exon 7 inclusion (**Figure 9A)**. We also tested CGCGCG in the above-mentioned *MAPT* minigene by inserting one copy at P32 in intron 10. However, the CG repeats markedly inhibited exon 10 splicing (**Figure 9B**), suggesting that the effect of CG repeats is context-dependent.

**Figure 9.**
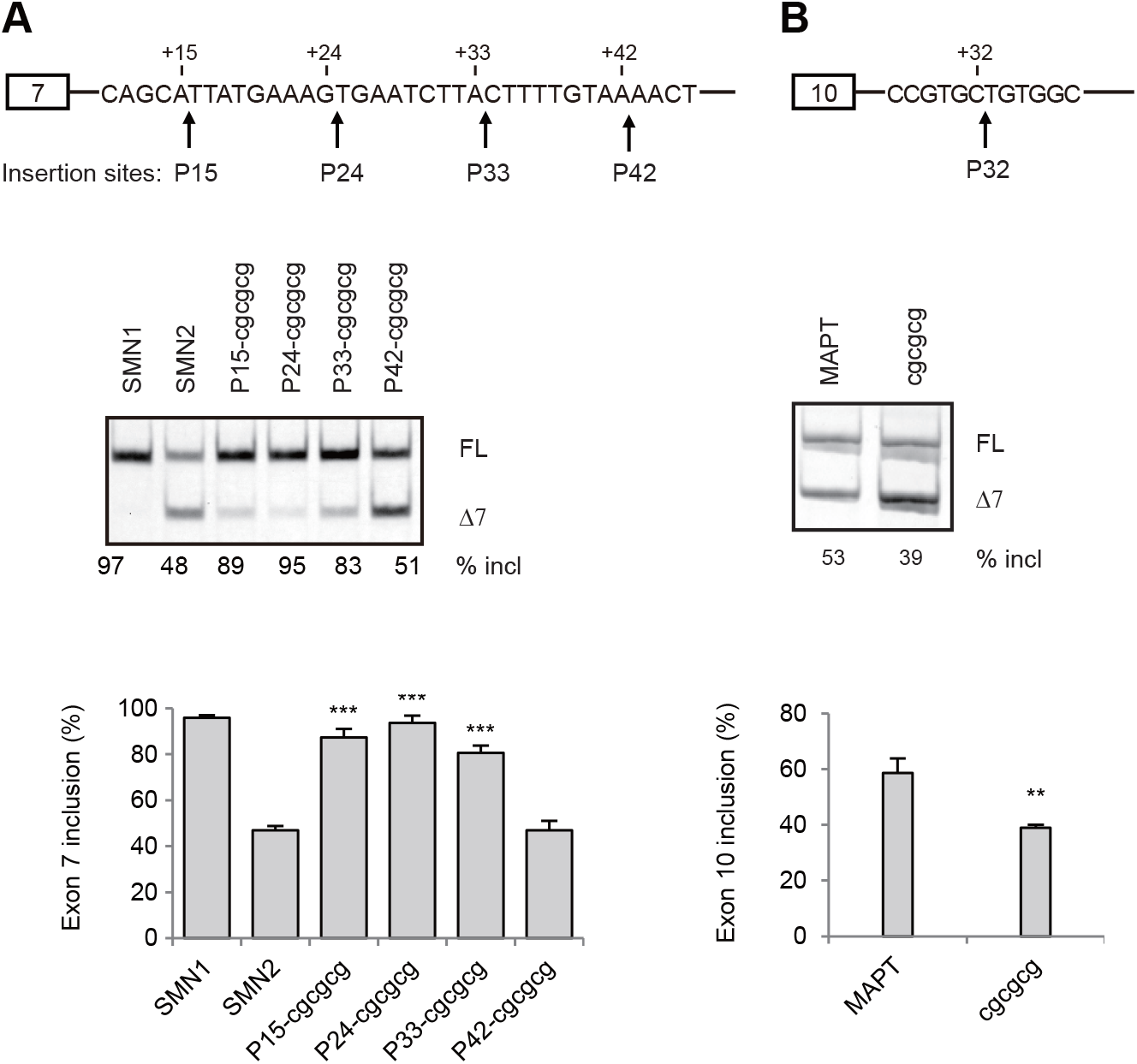
The effects of CG repeats on *SMN2* exon 7 and *MAPT* exon 10 splicing assayed in HEK293 cells. **(A)** We inserted CGCGCG at four sites: P15, P24, P33, and P42 of *SMN2* intron 7. Robust stimulatory effects are seen when the motif is closer to the 5′ splice site of intron 7 (P15, P24 and P33). **(B)** We also tested CGCGCG in the *MAPT* minigene by inserting it at P32 in intron 10, where an inhibitory effect is seen instead. Quantitative data are shown in histograms (*n* = 3). (**) *P* < 0.01, (***) *P* < 0.001 compared to WT *SMN2, SMN1 or MAPT*.

## DISCUSSION

Most RBPs recognize short sequence motifs to bind RNA and exert their functions (13). Although a cohort of SREs and their cognate proteins have been documented, it remains challenging to predict the patterns of sequence features that regulate alternative splicing. In the present study, we built a library of all possible pentameric sequences at a fixed site in intron 7 of an *SMN2* minigene by site-directed mutagenesis, and analyzed in duplicate the effects of all 1023 mutants, plus the wild type, on exon 7 splicing in HEK293 cells. Using MSEA, we found that 96 2-4-mer motifs are enriched in pentamers that robustly promote or repress exon 7 splicing. Based on their sequence features, these motifs were grouped into four stimulatory classes: U-rich, C-rich, CG-core and RBFOX-related, as well as four inhibitory classes: AG-core, UG-type, GCWCC-type, and GAC-core. The splicing effects of nearly all the top 50 most-stimulatory and most-inhibitory mutants are attributable to these motif classes (**Figure 2**). Therefore, our study revealed major classes of short intronic SRE motifs that potently affect alternative splicing. Three motif classes—CG-core, GCWCC, and GAC-core—are novel, and will require further investigation to understand their underlying mechanisms and characteristics in splicing regulation.

Systematic high-throughput screens of SREs have been reported in several studies using various strategies (21–26). A shared feature is that they all used random sequences coupled with RNA sequencing or selection by enrichment tools, such as GFP, as opposed to a complete sequence library being analyzed one by one—the approach we used here. Our study presents a full picture of splicing patterns for all pentameric sequences with respect to their effects on *SMN2* exon 7 splicing. Although different alternative splicing events may have distinct features, the general principles governing RNA splicing are widely applicable. Therefore, our results markedly improve our understanding of splicing regulation mediated by 5-nt or shorter sequences.

The length of random sequences used to generate a library in previous large-scale studies was 10 to 25 nt; analysis of short sequences, such as pentamers or hexamers, enriched within the longer sequence was key to identify functional motifs. We compared our data to those of prior studies. Interestingly, the strongest ISEs (CUUCUU, UAUUUU, UUGUUC, UCUUAU and UCUUAC) identified in a large-scale screen in HEK293 cells with 25-nt random sequences inserted downstream of alternative 5’ splice sites in an artificial citrine-based reporter by Rosenberg et al. are all U-rich motifs (22), which is highly consistent with our finding that U-rich sequences are the predominant class of stimulatory motifs. In fact, the top two hexamers CUUCUU and UAUUUU, which are equivalent to UUCUU and UAUUU, respectively, with the flanking C or U being included, are two of the strongest stimulatory pentamers in our library (**Figure 2**).

Wang et al. identified G-rich sequences as a group of the strongest ISEs in a GFP-based random-decamer screen in HEK293T cells (24), which is in broad agreement with our observation that G-rich sequences (4 and more consecutive Gs) markedly promote exon 7 splicing, though G-rich motifs were not among the strongest ISEs in the present study.

One of our salient findings is that CG-core motifs promote exon 7 splicing. Although this class of ISEs was not previously reported, we found that 8/10 top 6-mer ESEs identified by Rosenberg et al. harbor at least 1 CG repeat, and half of these have 2 CG repeats (22), suggesting that CG-core motifs may be common splicing enhancers present in both exons and introns, though they may be inhibitory in some contexts, like MAPT. Of note, the CG dinucleotide is under-represented in vertebrate genomes (40).

Two studies specifically explored intronic SREs with a 10- or 15-nt random sequence library in a GFP-fused *SMN1* minigene and a GFP-based reporter, respectively, in HEK293 cells, and a majority of their enriched 4-6-nt motifs comprise the dinucleotide AG (21,24), consistent with our data. Intronic SRE motifs identified by Culler et al. also include UG-rich and GACC (21), which highly resemble the UG-type and GAC-core motifs in our study.

On the other hand, we noted considerable discrepancies between our data and those in previous studies that relied on random-sequence pools. For example, in line with the established notion that the RBFOX-binding sequence UGCAUG is a strong ISE when placed downstream of the 5′ splice site of an alternative cassette exon (36), we found that UGCAUG, as seen with multiple (UGC)AUGNN mutants, robustly promotes exon 7 inclusion (**Figure 2**). However, UGCAUG was not identified in the previous high-throughput studies. Another example is the C-rich motif class, which we characterized as a main class of ISEs in the present study, but was likewise not found in the previous high-throughput studies. In addition, the two potent inhibitory pentamers GCACC and GCUCC we observed, were not detected as winner ISS motifs in prior random-sequence screens, despite the enrichment of weaker elements, CCUCC or CUCC, in two studies (21,25).

These discrepancies could be due to the different gene contexts, different cell types or conditions used for analysis, or effects of potential RNA secondary structures that affect splice-site recognition. Another key issue that was previously overlooked is the length of the random sequences. Based on previous studies, the binding sites of most RBPs are ~5 nt. In light of our observations, motifs as short as 2 nt, such as CG and AG, can determine whether a pentamer is stimulatory or inhibitory, and many 3-6 nt motifs, such as UCG, AGG, UAG, CAG, GAC, UGG, GGU, YCGY, UAGG, CAGG, UGGU, CUCC, GCACC, U-rich, C-rich, UA-rich, poly-G and UGCAUG, are potent SREs. When the length of the library sequences is 10 nt or longer, it is inevitable that each sequence harbors multiple motifs, giving rise to complex consequences. Some authentic strong enhancer motifs may be missed, due to silencer motifs being next to or overlapping them, and vice versa. The worst-case scenario is that an enhancer may be mistaken for a silencer, due to the presence of a nearby strong silencer, which, we believe, is relatively common. For example, UGCAUG itself is a known strong enhancer, and UGAC represents a strong silencer in our study; when both motifs overlap, as seen in mutant (UGC)AUGAC in our library, the silencer motif is dominant, and the net effect is moderately inhibitory (**Supplementary Table S1**).

Among the eight classes of stimulatory and inhibitory motifs enriched by MSEA, the U-rich, RBFOX-related, and hnRNP A1-binding AG-core motifs were already well established, with known mechanisms (9,14,35,41,42). Ji et al. revealed PCBP1 and PCBP2 as global splicing activators; co-depletion of the two proteins inhibits inclusion of cassette exons flanked by intronic C-rich motifs that are immediately adjacent to the 5′ and/or 3′ splice site (43). Though we have not investigated which RBP(s) may mediate the effect of the C-rich motifs in our mutant minigenes, it is reasonable to assume that their cognate proteins are PCBP1 and PCBP2. Zheng et al. uncovered UGGU as the core motif in an ESS that inhibits a 3′ splice site in a bovine papillomavirus (BPV) type 1 late transcript, but no cognate RBP was identified (44). Although UG-type motifs are highly enriched in strong inhibitory pentamers in our library, their roles in splicing regulation may be over-estimated, owing to interference with the intramolecular RNA structure that impairs U1 annealing (**Supplementary Figure S2**). GAC-core motifs represent a novel class of short ISSs. Though they have not been previously characterized, SELEX performed by Cavaloc et al. revealed that SRSF7 (formerly 9G8) binds with high affinity to GAC repeats (45). However, whether the silencing activity of the GAC-core motifs we analyzed is mediated by SRSF7—which has been characterized as a splicing activator, rather than a repressor—needs to be investigated.

Two notable findings in the present study are that the dinucleotide CG represents the core of the second most abundant class of stimulatory motifs, and that GCWCC-type motifs are potent ISSs. Whereas CG-core motifs or CG repeats will require further study to derive the consensus sequence, the winner sequences for GCWCC-type motifs are clearly GCACC and GCUCC. An early study by Zheng et al. delineated a so-called C-rich ESS sequence, GGCUCCCC, in BPV-1 pre-mRNA (44), which encompasses a GCUCC pentamer. It is intriguing that GCACC, despite being the second most inhibitory pentamer in our study, was not previously shown to regulate natural alternative splicing events. We believe that both CG-core and GCWCC motifs should play a widespread and important role in regulating alternative splicing, considering their potency in affecting splicing and expected relative frequency of occurrence of such short motifs in the human genome. Future studies should identify the RBPs that recognize these motifs, and their regulated alternative splicing targets, so as to gain insights into their significance in gene-expression regulation.

Although degeneracy is a typical feature for many RBPs, we observed that in many cases a single nucleotide change resulted in marked changes in exon 7 inclusion. The reason is that many core motifs are just 2-4 nt long, and a single nucleotide change can easily convert an enhancer into a silencer, for example, CG to AG, GGGG to AGGG and CCCC to CUCC, or vice versa. One surprising finding in the present study is the complexity of motifs comprising U and C. In the library, seven pentamers UUCUU, UCUUU, CUUUU, CCCUU, CUCUU, CCCCU and CCCUC are among the top 50 stimulatory ones, whereas four pentamers UCCUC, CUUCC, UUUCU and CCUCC are among the top 50 inhibitory ones. The sequence differences between them are quite subtle. A detailed study will be required to pinpoint the distinguishing sequence features and underlying mechanisms.

The pentamer library also provides a resource to better understand known motifs and their respective RBPs. For example, PTB is a strong splicing repressor whose known binding motifs are UCUU, UCUUC, and CUCUCU (46,47). We indeed found that overexpression of PTB in HEK293 cells strongly inhibits exon 7 splicing in the *SMN1* and *SMN2* minigenes (data not shown). Paradoxically, however, the pentamers NUCUU and UCUUN, as well as UCUCU and CUCUC (which form CUCUCU with flanking nucleotides) all potently stimulate exon 7 splicing. Both UCUU and UCUC are enriched ISE motifs by MSEA. Another example is the RBFOX protein-binding motif UGCAUG, a strong ISE, selected by the Position 1-dependent MSEA. Based on our data (**Supplementary Table S1**), exon 7 splicing is much improved when UGCAUG is followed by C or U compared to G or A. In contrast, when UGCAUG is followed by AC, its stimulatory effect disappears. These examples highlight the complexity of interactions between SREs and their cognate RBPs in regulating pre-mRNA splicing.

## Acknowledgment

Y.H. gratefully acknowledges the National Natural Science Foundation of China (grants 82073753, 81530035 and 81471298,). A.R.K. acknowledges support from NIH grant R37-GM42699.

## Data availability statement

All data generated or analyzed during this study are included in this published article and its supplementary information.

## Notes

\# Co-first authors.

### Competing Interest Statement

The authors have declared no competing interest.

