## Supplementary Figures 1 and 2, Tables S3-S9 for "Systematic characterization of short intronic splicing-regulatory elements"

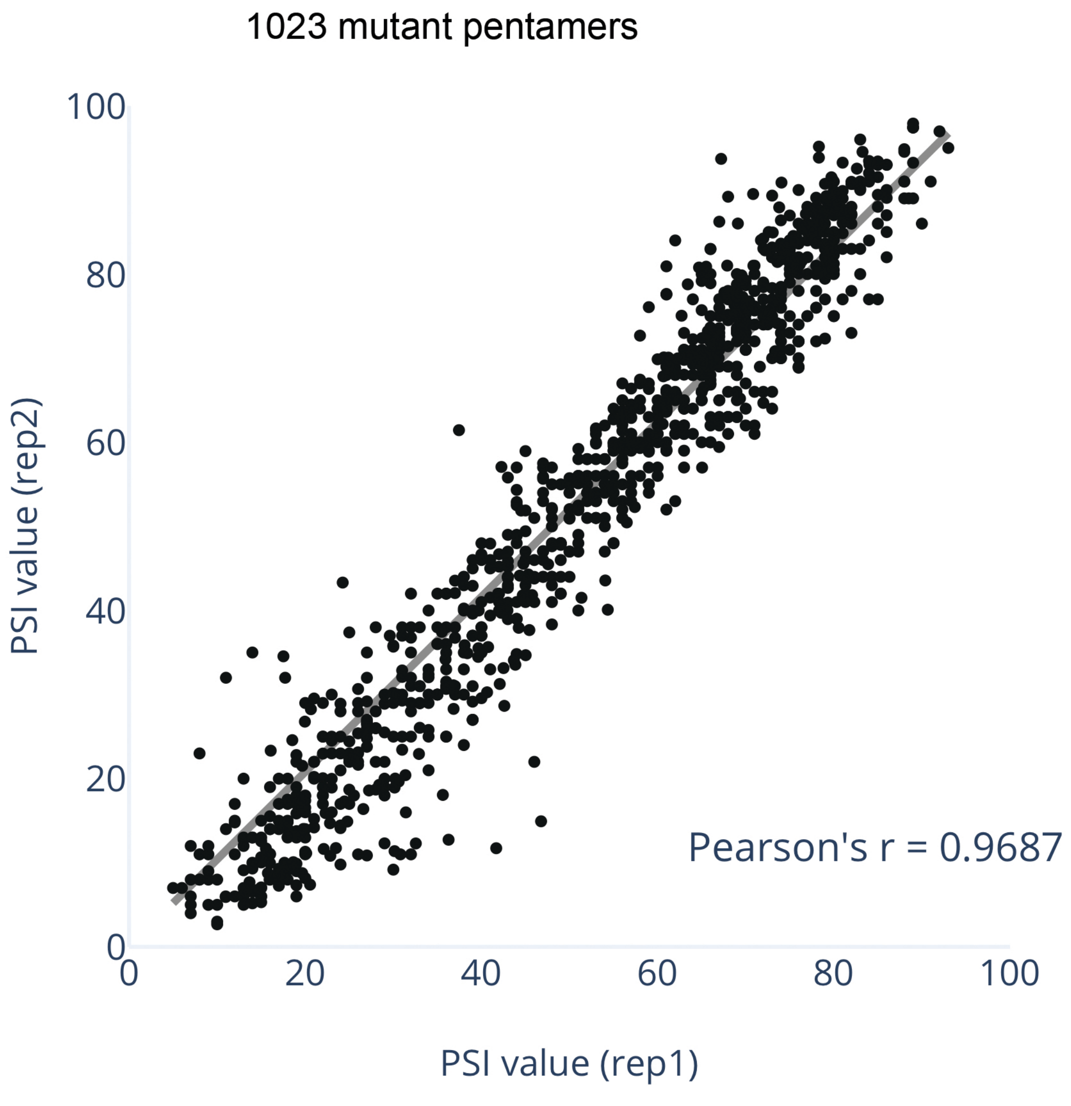


**Figure S1.** Scatterplot of PSI values of 1,023 mutant pentamers in two replicates. Each block dot represents on mutant pentamer. X- and Y-axes show the PSI value of *SMN2* exon 7 inclusion in two replicates (rep1 and rep2), respectively. Linear regression was used to calculate the $R^2$ value while intercept was set to zero (y = 1.0405x).

### Supplementary Figure S2

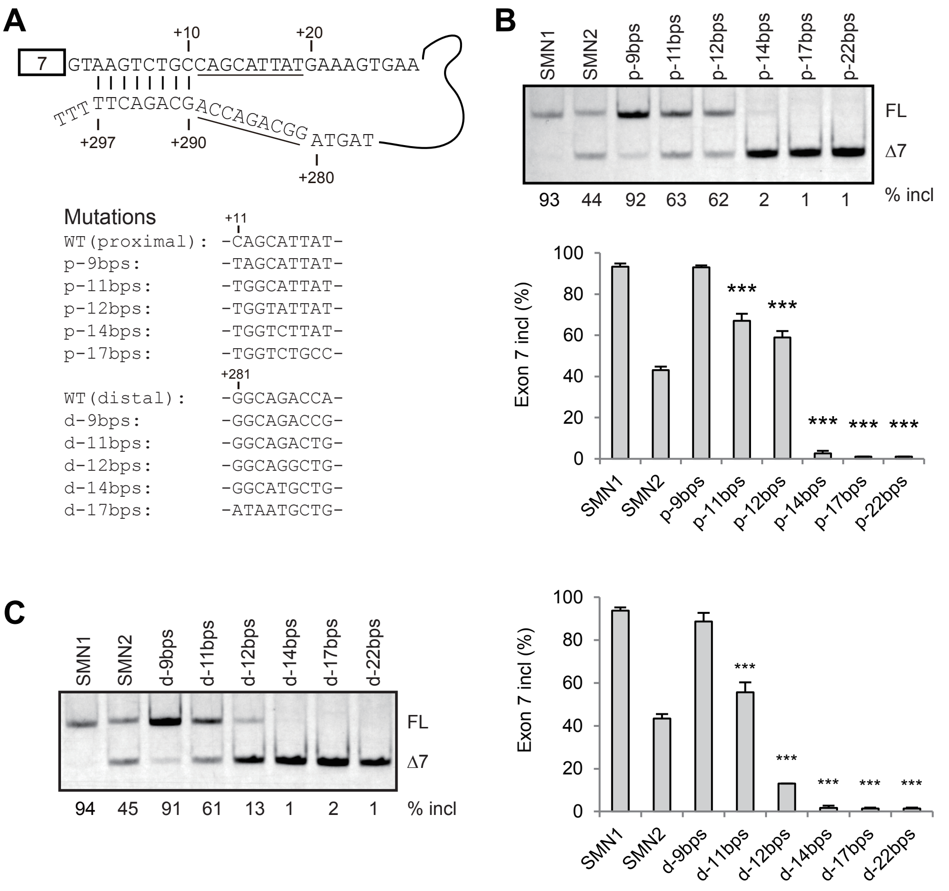


**Figure S2.** Strengthening the putative dsRNA in *SMN1* intron 7 inhibits exon 7 splicing. **A**. *SMN1* minigene mutants and their sequences were shown. Analysis of exon 7 splicing for mutations in the proximal (**B**) and distal (**C**) strands of the dsRNA structure in HEK293 cells by Cy5-labeled RT-PCR. Histagrams clearly show an inverse correlation between the length of the dsRNA stretch and the percentage of exon 7 inclusion (% incl). FL: full length; Δ7: exon 7-skipped isoform. *** $P$ < 0.001 compared to the WT *SMN2*.

**Supplementary Table S3**. All 40 pentamers comprised of U and/or A and their ΔPSI values. Note, pentamers with more Us usually had much higher ΔPSI values.

| Pentamer | ΔPSI | Pentamer | ΔPSI | Pentamer | ΔPSI | Pentamer | ΔPSI |
|---|---|---|---|---|---|---|---|
| AAAAA | 10.50 | AUAAA | 16.50 | UAAAA | 1.50 | UUAAA | 10.50 |
| AAAAU | 27.00 | AUAAU | 39.50 | UAAAU | 27.00 | UUAAU | 31.50 |
| AAAUA | 22.00 | AUAUA | 33.50 | UAAUA | 25.50 | UUAUA | 41.50 |
| AAAUU | 35.50 | AUAUU | 44.50 | UAAUU | 37.50 | UUAUU | 46.00 |
| AAUAA | 14.00 | AUUAA | 28.50 | UAUAA | 11.00 | UUUAA | 23.50 |
| AAUAU | 42.00 | AUUAU | 49.00 | UAUAU | 36.00 | UUUAU | 50.50 |
| AAUUA | 33.00 | AUUUA | 51.00 | UAUUA | 37.00 | UUUUA | 57.00 |
| AAUUU | 45.50 | AUUUU | 51.50 | UAUUU | 47.50 | UUUUU | 58.50 |

**Supplementary Table S4**. All 40 pentamers comprised of U and/or C and their ΔPSI values. While most of these pentamers strongly promoted *SMN2* exon 7 splicing, some others were very inhibitory.

| Pentamer | ΔPSI | Pentamer | ΔPSI | Pentamer | ΔPSI | Pentamer | ΔPSI |
|---|---|---|---|---|---|---|---|
| CCCCC | 41.50 | CUCCC | 8.50 | UCCCC | 12.00 | UUCCC | 25.50 |
| CCCCU | 46.00 | CUCCU | -3.50 | UCCCU | 24.02 | UUCCU | 38.05 |
| CCCUC | 46.00 | CUCUC | 41.00 | UCCUC | -35.91 | UUCUC | 40.50 |
| CCCUU | 48.50 | CUCUU | 48.00 | UCCUU | -24.50 | UUCUU | 58.50 |
| CCUCC | -28.50 | CUUCC | -30.50 | UCUCC | -3.50 | UUUCC | -12.50 |
| CCUCU | 39.23 | CUUCU | 21.78 | UCUCU | 38.97 | UUUCU | -30.34 |
| CCUUC | -11.50 | CUUUC | 31.39 | UCUUC | 42.27 | UUUUC | 45.48 |
| CCUUU | 44.00 | CUUUU | 52.50 | UCUUU | 55.50 | UUUUU | 58.50 |

**Supplementary Table S5**. List of all 16 pentamers containing UGGY motifs and their ΔPSI values.

| Pentamer | ΔPSI | Pentamer | ΔPSI |
|---|---|---|---|
| UGGUA | -28.50 | UGGCA | -26.00 |
| UGGUC | -42.66 | UGGCC | -31.00 |
| UGGUG | -32.50 | UGGCG | -26.47 |
| UGGUU | -24.50 | UGGCU | -27.48 |
| AUGGU* | 14.50 | AUGGC* | -2.35 |
| CUGGU | -20.00 | CUGGC | -18.63 |
| GUGGU | -15.00 | GUGGC | -20.40 |
| UUGGU | -19.00 | UUGGC | -14.80 |

* They form RBFOX-binding motifs (UGCAUG) with the upstream trinucleotide UGC.

**Supplementary Table S6**. List of all 16 NUGGN pentamers (N = A, C, G or U) and their ΔPSI values.

| Pentamer | ΔPSI | Pentamer | ΔPSI | Pentamer | ΔPSI | Pentamer | ΔPSI |
|---|---|---|---|---|---|---|---|
| AUGGA | 39.50 | CUGGA | -10.50 | GUGGA | 19.50 | UUGGA | 0.00 |
| AUGGC | -2.35 | CUGGC | -18.63 | GUGGC | -20.40 | UUGGC | -14.80 |
| AUGGG | 32.41 | CUGGG | -4.84 | GUGGG | 34.01 | UUGGG | 0.61 |
| AUGGU | 14.5 | CUGGU | -20.00 | GUGGU | -15.00 | UUGGU | -19.00 |

**Supplementary Table S7**. List of all 40 GGG-containing pentamers. 30 pentamers have a ΔPSI > 0; 18 have a ΔPSI > 17. Pentamers containing GGGG are shaded.

| Pentamer | ΔPSI | Pentamer | ΔPSI | Pentamer | ΔPSI | Pentamer | ΔPSI |
|---|---|---|---|---|---|---|---|
| GGGAA | 21.50 | GGGGG | 39.02 | CGGGA | 9.50 | AUGGG | 32.41 |
| GGGAC | 2.26 | GGGGU | 26.50 | CGGGC | 5.30 | CAGGG | -23.89 |
| GGGAG | 1.48 | GGGUA | 30.50 | CGGGG | 27.98 | CCGGG | 27.95 |
| GGGAU | 16.00 | GGGUC | 4.17 | CGGGU | 18.50 | CUGGG | -4.84 |
| GGGCA | 21.50 | GGGUG | 24.95 | UGGGA | 2.00 | GAGGG | -13.96 |
| GGGCC | 5.50 | GGGUU | 4.50 | UGGGC | -24.14 | GCGGG | 41.53 |
| GGGCG | 26.89 | AGGGA | -33.00 | UGGGG | 1.34 | GUGGG | 34.10 |
| GGGCU | 20.34 | AGGGC | -36.52 | UGGGU | -21.50 | UAGGG | -31.96 |
| GGGGA | 40.50 | AGGGG | -17.97 | AAGGG | 3.94 | UCGGG | 1.10 |
| GGGGC | 27.23 | AGGGU | -25.00 | ACGGG | 17.42 | UUGGG | 0.61 |

**Supplementary Table S8**. List of all 30 pentamers containing CGAC (CGACN and NCGAC) or creating CGAC with the upstream nucleotide C (GACNN) and their ΔPSI values.

| Pentamer | ΔPSI | Pentamer | ΔPSI | Pentamer | ΔPSI | Pentamer | ΔPSI |
|---|---|---|---|---|---|---|---|
| GACAA | 11.00 | GACCG | 39.75 | GACUA | 24.50 | CGACG | 24.44 |
| GACAC | 26.09 | GACCU | 37.62 | GACUC | 32.06 | CGACU | 14.11 |
| GACAG | -16.15 | GACGA | 40.00 | GACUG | 32.36 | ACGAC | 4.41 |
| GACAU | 29.00 | GACGC | 40.36 | GACUU | 43.50 | CCGAC | 2.64 |
| GACCA | 26.00 | GACGG | 34.19 | CGACA | 13.00 | GCGAC | -17.90 |
| GACCC | 17.50 | GACGU | 40.00 | CGACC | 7.50 | UCGAC | -16.81 |

**Supplementary Table S9**. Analysis of all pentamers containing DGAC.

| Pentamer | ΔPSI | Pentamer | ΔPSI | Pentamer | ΔPSI | Pentamer | ΔPSI |
|---|---|---|---|---|---|---|---|
| AGACA | -12.00 | GGACG | -20.65 | AAGAC | -28.06 | GGGAC | 2.26 |
| AGACC | -16.50 | GGACU | -20.00 | CAGAC | -30.78 | UGGAC | -36.34 |
| AGACG | -14.15 | UGACA | -5.50 | GAGAC | -33.39 | AUGAC | -23.24 |
| AGACU | -12.78 | UGACC | -1.50 | UAGAC | -32.34 | CUGAC | -21.67 |
| GGACA | -18.00 | UGACG | -17.09 | AGGAC | -28.11 | GUGAC | -33.02 |
| GGACC | -6.00 | UGACU | -9.01 | CGGAC | 16.24 | UUGAC | -16.00 |
